# Ten Years of Using Key Characteristics of Human Carcinogens to Organize and Evaluate Mechanistic Evidence in IARC Monographs on the Identification of Carcinogenic Hazards to Humans: Patterns and Associations

**DOI:** 10.1101/2023.07.11.548354

**Authors:** Ivan Rusyn, Fred A. Wright

**Affiliations:** Department of Veterinary Pharmacology and Physiology, Texas A&M University, College Station, TX, USA; Departments of Statistics and Biological Sciences, and Bioinformatics Research Center, North Carolina State University, Raleigh, NC, USA

## Abstract

Systematic review and evaluation of the mechanistic evidence only recently been instituted in cancer hazard identification step of decision-making. One example of organizing and evaluating mechanistic evidence is the Key Characteristics approach of the International Agency for Research on Cancer (IARC) Monographs on the Identification of Carcinogenic Hazards to Humans. The Key Characteristics of Human Carcinogens were proposed almost 10 years ago and have been used in every IARC Monograph since 2015. We investigated the patterns and associations in the use of Key Characteristics by the independent expert Working Groups. We examined 19 Monographs (2015-2022) that evaluated 73 agents. We extracted information on the conclusions by each Working Group on the strength of evidence for agent-Key Characteristic combinations, data types that were available for decisions, and the role mechanistic data played in the final cancer hazard classification. We conducted both descriptive and association analyses within and across data types. We found that IARC Working Groups were cautious when evaluating mechanistic evidence: for only ∼13% of the agents was strong evidence assigned for any Key Characteristic. Genotoxicity and cell proliferation were most data-rich, while little evidence was available for DNA repair and immortalization Key Characteristics. Analysis of the associations among Key Characteristics revealed that only chemical’s metabolic activation was significantly co-occurring with genotoxicity and cell proliferation/death. Evidence from exposed humans was limited, while mechanistic evidence from rodent studies *in vivo* was often available. Only genotoxicity and cell proliferation/death were strongly associated with decisions on whether mechanistic data was impactful on the final cancer hazard classification. The practice of using the Key Characteristics approach is now well-established at IARC Monographs and other government agencies and the analyses presented herein will inform the future use of mechanistic evidence in regulatory decision-making.

## Introduction

The International Agency for Research on Cancer (IARC) Monographs on the Identification of Carcinogenic Hazards to Humans has been in existence for over 50 years (Pearce et al. 2015). Since December of 1971, working groups of international experts that typically meet three times each year produced classifications for over 1,000 agents (chemicals, occupational and environmental exposures, physical and biological agents, personal habits, and dietary factors). The evaluations follow a rigorous and transparent process detailed in the Preamble to the Monographs (IARC Monographs Programme 2019). The most recent update to the Preamble was made in 2019 (Samet et al. 2020). IARC evaluations classify agents into four groups: 1 – carcinogenic to humans (126 agents, as of May 2023), 2A – probably carcinogenic to humans (94), 2B – possibly carcinogenic to humans (322), and 3 – not classifiable as to its carcinogenicity to humans (500).

The IARC Monographs process is designed to evaluate strength of the available evidence on cancer hazard (i.e., is the agent capable of causing cancer?) using studies in humans (e.g., cancer epidemiology), animals (e.g., chronic guideline tests in rodents), as well as mechanistic and other evidence. The latter has been traditionally used as a follow-up to a cancer hazard classification based on studies of cancer in humans and animals – for consideration of the potential “mechanistic” upgrade (e.g., a chemical belongs to a class of known human carcinogens, or strong evidence a known cancer mechanism operates in humans exposed to an agent) or downgrade (e.g., strong evidence that animal cancer findings are not relevant to humans) (IARC Monographs Programme 2006). According to the current IARC Monographs Preamble (IARC Monographs Programme 2019), mechanistic data can be used in combination with limited human or sufficient animal evidence of cancer hazard, or can be used by itself to classify an agent as possibly carcinogenic (Group 2B). These changes to the Preamble formalized several important developments that were pioneered in the IARC Monographs program in the areas such as systematic review using standardized workflows (Shapiro et al. 2018) and greater emphasis on mechanistic evidence through the Key Characteristics of Carcinogens approach (Smith et al. 2016).

Key Characteristics of Carcinogens were defined through several expert workshops in 2012 that were convened by IARC to evaluate tumor concordance (Krewski et al. 2019b) and mechanisms of carcinogenesis (Birkett et al. 2019) information for agents previously classified as Group 1. Contemporaneous publications demonstrating that cancer cells (Hanahan and Weinberg 2011) and agents that cause cancer (Guyton et al. 2009; Kushman et al. 2013) possess a finite set of common characteristics, combined with the systematic analysis of diverse known human carcinogens evaluated by IARC Monographs in the previous 40 years (Al-Zoughool et al. 2019), opened a path to define properties that carcinogenic agents commonly show. Such an approach to identify, organize, and summarize results from often voluminous mechanistic studies aimed to introduce objectivity and promote structured weight-of-evidence expert judgement by diverse panels of experts who participate in the IARC Monographs Working Groups (Smith et al. 2016). Key Characteristics have been in use at IARC Monographs since March 2015 (Guyton et al. 2015), first based on the Monograph-specific instructions (IARC Monographs Programme 2021) provided to the members of the Working Group and others participating in each meeting, and more recently as detailed in the IARC Monographs Preamble (IARC Monographs Programme 2019). Use of high-throughput *in vitro* toxicity screening data in cancer hazard evaluations by IARC Monographs was pioneered together with Key Characteristics (Chiu et al. 2018) and was among the first examples of these data being used in decision-making.

Indeed, the wide recognition of the greater role for mechanistic evidence in cancer hazard identification, as well as in other areas of regulatory decisions for industrial (U.S. EPA 2021, 2022), food-use (Alimohammadi et al. 2023) and drugs (Strauss et al. 2021), has been catalyzed by both the time and expense of human epidemiology and chronic animal studies, considerations of animal welfare, and advances in computational and *in vitro* models in toxicology (Andersen and Krewski 2010; Kavlock et al. 2018; Krewski et al. 2020; Sturla et al. 2014). In fact, there has been no shortage of the frameworks proposed to encourage decision-makers to embrace data from non-human/-animal studies (Anklam et al. 2022; Berggren et al. 2015; Hoffmann et al. 2022; Krewski et al. 2022). The traditional mode-of-action framework (Ashby et al. 1996), adopted by the International Programme on Chemical Safety (Meek et al. 2014) and United States Environmental Protection Agency (US EPA) (U.S. Environmental Protection Agency 2005), has been most widely used to consider human relevance of cancer findings in animals, and/or to justify selection of linear/non-linear dose response (Boobis et al. 2006; Boobis et al. 2008). More recently, several international expert groups (e.g., Organization for Economic Cooperation and Development, OECD) assembled *in silico* and *in vitro* test methods into so called Integrated Approaches to Testing and Assessment (IATA) using the concept of Adverse Outcome Pathways (AOPs), a framework for organizing data from different methods and across levels of biology to evaluate relationships between key events and adverse effects (Edwards et al. 2016; Patlewicz et al. 2014). Most advanced examples of such IATAs transforming into test guidelines are for skin sensitization (OECD 2016, 2021), and developmental neurotoxicity (OECD 2022). Additional case studies and advantages and limitations of this approach have been recently published (Bajard et al. 2023).

Still, while there are several “mechanistic constructs” that, when used in conjunction with evidence from mechanistic studies, can inform and potentially replace traditional data from human and animal studies (Meek and Wikoff 2023), it may take a decade or more before they impact specific regulatory decisions. It takes time to convince regulatory agencies that their processes should evolve, and chemical assessments may require decades to be finalized (National Research Council 2014). Conversely, the IARC Monographs process is typically more expedient, taking approximately one year from the announcement of the list of agents and a call for data and experts, to the expert panel meeting where the final evaluations are decided and announced through a short summary in Lancet Oncology journal. It may take another year or more for the final Monograph to appear in press. It is not surprising, therefore, that a relatively short time (∼2 years) has passed from the conceptualization of Key Characteristics to their finalization and application in the first Monograph (Guyton et al. 2015). Since 2015, nineteen IARC Monographs (volumes 112-130) that were available in full text included a total of 73 agents, and all of these underwent mechanistic evidence evaluation using the Key Characteristics approach (Supplemental Table 1). While a summary of the first two years of application of Key Characteristics by IARC Monographs has been published (Guyton et al. 2018a), a far more diverse and larger set of agents is available today, ten years since the Key Characteristics approach was formalized. Over a hundred experts participated in these evaluations, and beyond a common set of instructions these experts used their independent weight-of-evidence considerations to exercise expert judgement on both strength and cohesion in the mechanistic evidence for each agent and Key Characteristic. This study extracted both Working Group conclusions from each IARC Monograph (Supplemental Table 1), as well as evaluated the consistency of mechanistic evidence from different model systems to determine both patterns and associations within and between evidence types, Key Characteristics, and their ultimate use for final classifications.

## Methods

### Extraction of the information from IARC Monographs

Full text PDF files for each IARC Monograph from Volume 112 to Volume 130 were downloaded from the IARC Monographs website https://monographs.iarc.who.int/monographs-available/ (see Supplemental Table 1 for hyperlinks). Information from Chapters 4 (Mechanistic and Other Relevant Data), 5.4 (Summary of Data Reported: Mechanistic and other relevant data), and 6.3 (Evaluation: Overall evaluation) was coded using standardized terminology and placed into spreadsheets (Supplemental Table 2). For each Monograph, each agent’s name, and its final classification and conclusions regarding the strengths of human and animal evidence (sufficient, limited, or inadequate), along with an indication of whether mechanistic data may have played a role in the final classification upgrade, supportive, or not used) were extracted from chapter 6.3. These conclusions were transcribed exactly as made by the Working Groups.

In addition, 8 categories were created to describe a “model system” for which mechanistic data may have been available during the evaluations. These included categories of “exposed humans” (studies on specimens collected from human subjects with documented exposures), “human *in vitro*” (studies in primary or other cell types of human origin that were exposed to an agent outside of the human body), “mammalian *in vivo*” (studies on specimens collected from live animals exposed to an agent), “mammalian *in vitro*” (studies in primary or other cell types of animal origin that were exposed to an agent outside of the animal body), “other *in vivo*” (studies on specimens collected from live non-mammalian vertebrates that were exposed to an agent), “other *in vitro*” (studies in either primary or other cell types collected from non-mammalian vertebrates, or in lower organisms (worms, yeast or prokaryotes) that were exposed to an agent), “ToxCast data” (data from assays accessible through US EPA Dashboard (Williams et al. 2017)), and “ToxRefDB data” (data from chronic animal studies from the US EPA ToxRef database (Martin et al. 2009)). These categories are largely concordant with those suggested in the IARC Monographs Preamble (IARC Monographs Programme 2019), but modified somewhat to enable further analyses (see below). For each agent, Key Characteristic, and model type, the information in Chapter 4 of each Monograph was examined to determine whether studies cited in the Monograph were supportive (“yes”, e.g., several studies were summarized indicating that a Key Characteristic was operational), “equivocal” (e.g., there were few studies, or among several studies summarized the evidence was pointing both ways), opposing (“no”, e.g. several studies were available that tested for measures of a Key Characteristic and found no evidence for it being involved), or not available (blank). The readers should note that these determinations were made by the authors of this manuscript, not IARC Monographs Working Group members, using their best professional judgement and experience with similar evaluations.

Finally, the “overall strength” of evidence for each Key Characteristic was transcribed from Chapter 5.3 of each Monograph. The terminology that was used to determine strength of mechanistic evidence for each Key Characteristic has evolved over time between Volumes 112 and 130. In Volumes 112-124, the Working Groups most often used strong/moderate/weak terms when data was available to decide for a particular Key Characteristic. From Volume 125, the terminology changed to consistent/suggestive/conflicting/inconsistent. The exact terms used by each Working Group were recorded (Supplemental Table 2); standardization of the terminology is detailed below.

### Data transformation

To enable data analyses, data in Supplemental Table 2 were modified in two ways. First, the “overall strength” descriptors were standardized as follows – (i) *strong*, *moderate*, and *weak* descriptors were retained as stated; (ii) *consistent* descriptors were substituted for *strong*, *suggestive* for *moderate*, and *conflicting* or *inconsistent* for *weak*. Second, data were transposed to retain agent name, final classification, mechanistic data role, and overall strength for each Key Characteristic. The original and standardized terms for “overall strength” are presented as Supplemental Tables 3 and 4, respectively. The standardized terminology (strong, moderate, or weak) was used in the analyses detailed below.

### Data analyses (descriptive statistics)

Column- and row-wise analyses using information in Supplemental Tables 2 and 3 were used for visualization and significance testing. Ordinary one-way ANOVA analysis was used to compare the mean of each column with the mean of every other column, after determining that the means were large enough that ANOVA assumptions were appropriate or conservative. The results were corrected for multiple comparisons using the Tukey method. P-values adjusted for multiple comparisons (p<0.05) were deemed significant. These analyses were conducted using GraphPad Prism v.9.4.1. (San Diego, CA).

### Data analyses (associations)

For analysis type (I) (see Figure 1), 45 (10 choose 2) pairwise comparisons of key characteristics for the Overall Summary were performed using Armitage’s trend test (two-sided). To ensure that p-values were robust for small sample sizes, Fisher’s exact test was used to filter out instances in which the exact test *p-*value was >10 fold larger than the trend p-value. The same approach was used for analysis type (II) (ToxCast data). For analysis types (III) and (IV), ToxCast determinations were similarly analyzed for associations using Fisher’s exact test only, due to small cell counts. The Benjamini-Hochberg approach (Benjamini and Hochberg 1995) was used to control the false discovery rate using the R p.adjust function, which automatically adapts to instances in which some pair-wise comparisons were not possible, with false discovery *q*<0.05 declared significant. Analyses were performed using R (v4.1.2).

**Figure 1.**
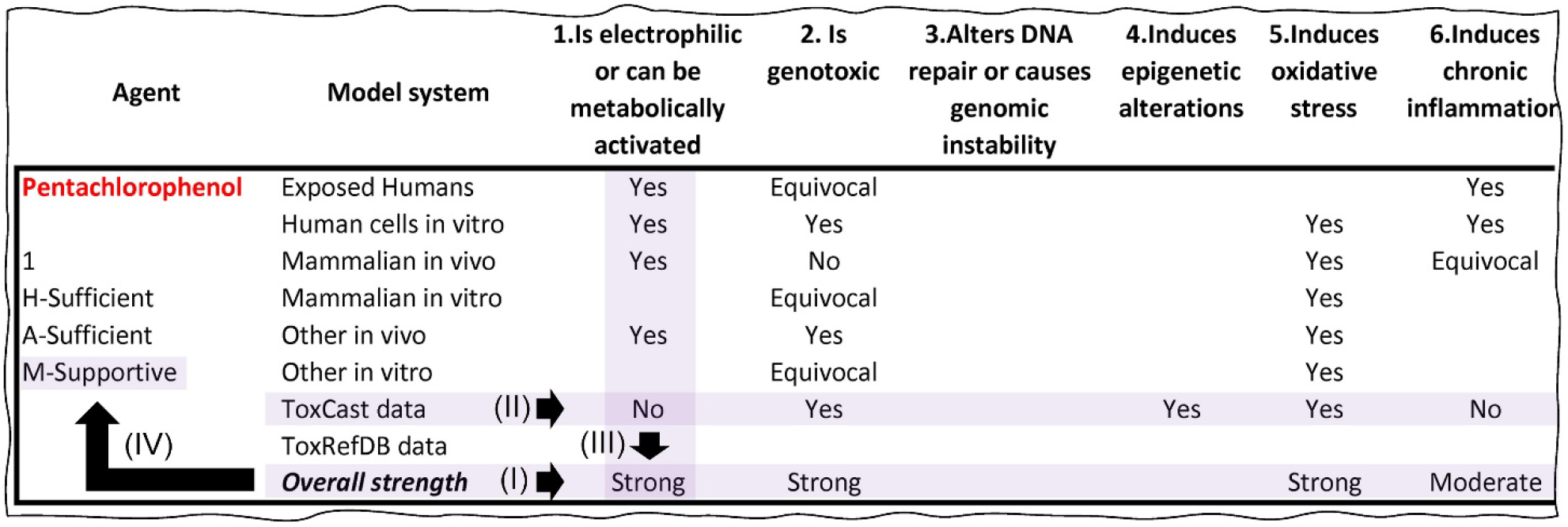
Illustration of a portion of the IARC Monographs dataset used in this study (Supplemental Table 2), for agent Pentachlorophenol and the first six Key Characteristics. Association analysis type (I) compares the Overall Strength values across all pairs of Key Characteristics. Analysis type (II) similarly compares the Key Characteristics for ToxCast data. Analysis Type (III) compares the ToxCast data as associated with Overall Strength within each Key Characteristic, and analysis type (IV) compares Overall Strength with how the mechanistic data weighed on the overall Cancer Hazard Classification.

## Results

### Frequency of use of Key Characteristics by IARC Monographs Working Groups

Previous retrospective analysis of the mechanistic data from animals and humans for 86 known human carcinogens (Group 1) identified by IARC Monographs Working Groups through volume 106 (Al-Zoughool et al. 2019) concluded that sufficient evidence was present for between 1 and 9 key characteristics (4.1±2.1 on average) per agent (Krewski et al. 2019a). This analysis was conducted by a group of experts, not IARC Monograph Working Group members, and did not include formal determination of the strength (i.e., strong, moderate, or weak) of evidence for each characteristic. There was also no key characteristics-based re-analysis of the agents previously classified by IARC as Group 2A, 2B, or 3, thus precluding any conclusions as to whether the mechanistic database was substantially different between agents in Group 1 and those that did not have sufficient evidence of cancer in humans (or other attributes that would have upgraded them to Group 1). Therefore, formalization of the process of key characteristics-based evaluations of the mechanistic and other evidence by IARC Working Groups starting from Monograph 112 (Guyton et al. 2015) allows for the analysis of trends in the application of this mechanistic construct by independent groups of experts, across a wide range of agents and ultimate classifications.

Since Monograph 112, determinations of the strength of evidence for each of 10 key characteristics were performed for 73 agents (67 chemicals, 4 dietary/lifestyle factors and 2 occupations). Five of these (∼7%) were classified as Group1, 23 (∼32%) as Group 2A, 41 (∼56%) as Group 2B, and 4 (∼5%) as Group 3. Of all possible pairwise combinations of agents and key characteristics, the Working Groups concluded for ∼2/3 of the instances that there was insufficient data (Figure 2A). The range of key characteristics with no data was between 2 and 10 and a median of 7, demonstrating the overall paucity of the mechanistic evidence for most agents that were suspected to pose human cancer hazard. Of the remaining ∼1/3 of determinations, most key characteristics were classified as strong or moderate (∼13% each); however, on average, only one key characteristic per agent was deemed as strong, moderate, or weak. In fact, there was a strongly significant (*p*<0.001) difference between the number of key characteristics per agents with no data as compared to other groups, but no difference between strong, moderate, or weak categories (Figure 2B). When the data was examined for each key characteristic (Figure 2C), it was evident that “2.Is genotoxic” had the most agents with strong data, and fewest instances where none was available. Other notable key characteristics with most complete datasets, but still for only ∼50% or fewer agents, were “10.Alters cell proliferation, cell death or nutrient supply,” “5.Oxidative stress,” and “1.Is electrophilic or can be metabolically activated.” Interestingly, there is a perfect overlap between the relative rank (by frequency of their determination across agents) of these four key characteristics in the analysis presented herein and that of Krewski et al (2019a). The remaining Key Characteristics had fewer determinations; fewest agents with data were for “3.Alters DNA repair or causes genomic instability,” “4.Induces epigenetic alterations,” and “9.Causes immortalization.”

**Figure 2.**
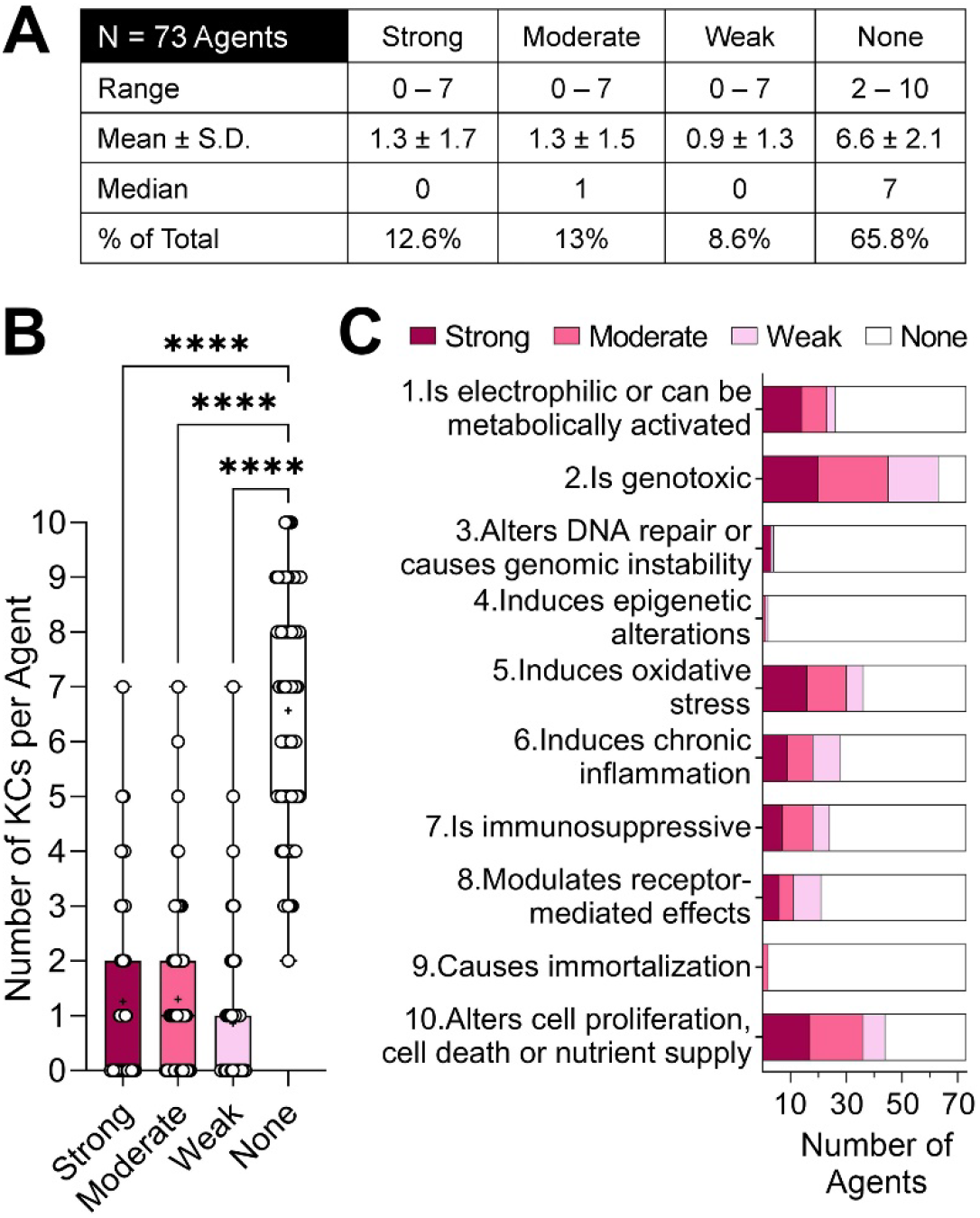
Summaries of Key Characteristics use in IARC Monographs 112-130. (A) Summaries of numbers of Key Characteristics with various strength of evidence designations. (B) Comparison of numbers of Key Characteristics per agent by strength of evidence. (C) Inverted stacked barplots of strength of evidence for each Key Characteristic.

Figure 3 presents the results of the analysis of key characteristic/agent combinations separately for the cancer hazard classification and strength of evidence. While only 5 agents among 73 evaluated by IARC across 19 Monographs since Key Characteristics started being formally used by the Working Groups were classified as Group 1, Key Characteristics with strong evidence were most typical (Figure 3A); however, among these 5 agents there was a wide range from 0 to 5 strong Key Characteristics per agent. Some additional Key Characteristics for these agents were deemed to be moderate or weak. For agents classified as Group 2A or 2B, no pattern was evident by the strength of evidence; several agents in these groups had as many strong Key Characteristics determinations as did Group 1 agents. When the data were separated by strength of evidence and cancer hazard classes were compared (Figure 3B), significant differences were observed. On average, agents classified as Group 1 and Group 2A had more strong Key Characteristics than those classified as Group 2B, but there was no difference between the former. There was also a general trend of erosion in the strength of evidence for each Key Characteristic from Group 1 to Group 3 (Figure 3C). Two notable exceptions to the trend were “10.Alters cell proliferation, cell death or nutrient supply” and “8.Modulates receptor-mediated effects” Key Characteristics for which agents classified as Group 2A were as many, or even more, as those in Group 1. It is noteworthy that several agents with strong Key Characteristic determinations were classified in Group 2B – a mere presence of a strong determination for any Key Characteristic was not an indication that an agent would be upgraded to a higher class; however, no agent classified as Group 3 had a strong key characteristic.

**Figure 3.**
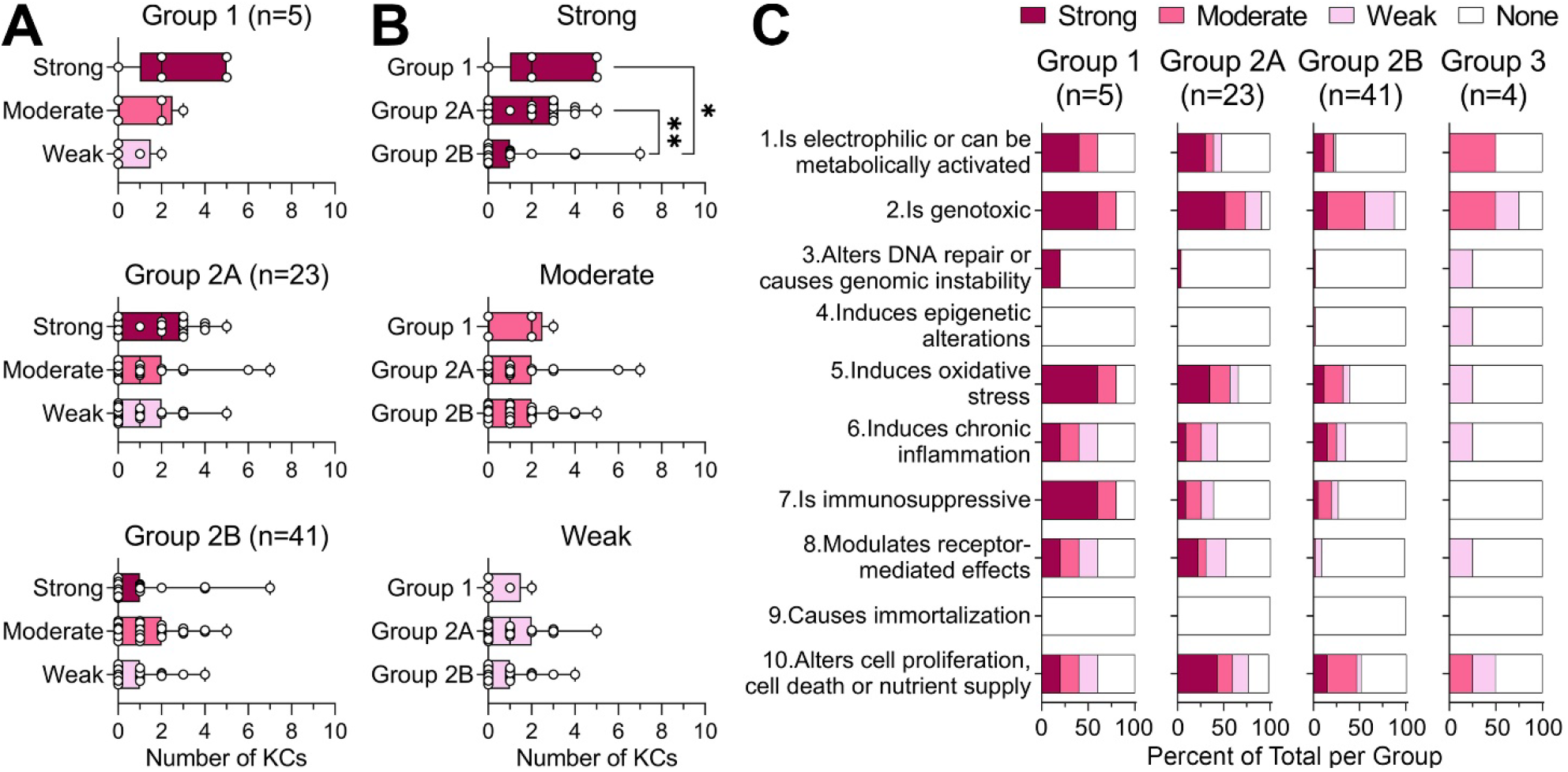
Cancer Hazard Classification and strength of evidence for Key Characteristics used in IARC Monographs 112-130. (A) Numbers of agents for each strength of evidence value for each Cancer Hazard Classification. (B) Numbers of agents for each Cancer Hazard Classification displayed for each strength of evidence value for each strength of evidence value. (C) Inverted stacked barplots displaying the strength of evidence for each Key Characteristic, with columns displaying the Cancer Hazard Classification.

It was next determined whether the strength of evidence for individual Key Characteristics, or their number, were different among agents for which the Working Groups deemed mechanistic evidence to be either sufficiently informative for an upgrade (9 agents), supportive of the classification determined based on human and/or animal data alone (15 agents), or not useable (49 agents). For agents that were upgraded using mechanistic data (Figure 4A, top) a significantly greater number of strong Key Characteristics, with a median of 3, was determined as compared to those that were moderate or weak. For agents where mechanistic data were supportive (Figure 4A, middle), the same was true but only as compared to those that were determined as weak. In cases when mechanistic data was not used (Figure 4A, bottom), significantly fewer strong Key Characteristics were determined as compared to those that were moderate. Overall, a clear significant trend was observed for Key Characteristics that were strong (Figure 4B) – both upgrade and supportive determinations had significantly more Key Characteristics. It should be noted that there was a wide range in the number of Key Characteristics in each category. With regards to the strength of evidence for each Key Characteristic (Figure 4C), three were deemed strong in cases of mechanistic upgrades – “2.Is genotoxic,” “10.Alters cell proliferation, cell death or nutrient supply,” and “1.Is electrophilic or can be metabolically activated.” The “5.Induces oxidative stress” Key Characteristic was most used in cases where mechanistic evidence was deemed to be supportive of the classification based on human and/or animal data alone.

**Figure 4.**
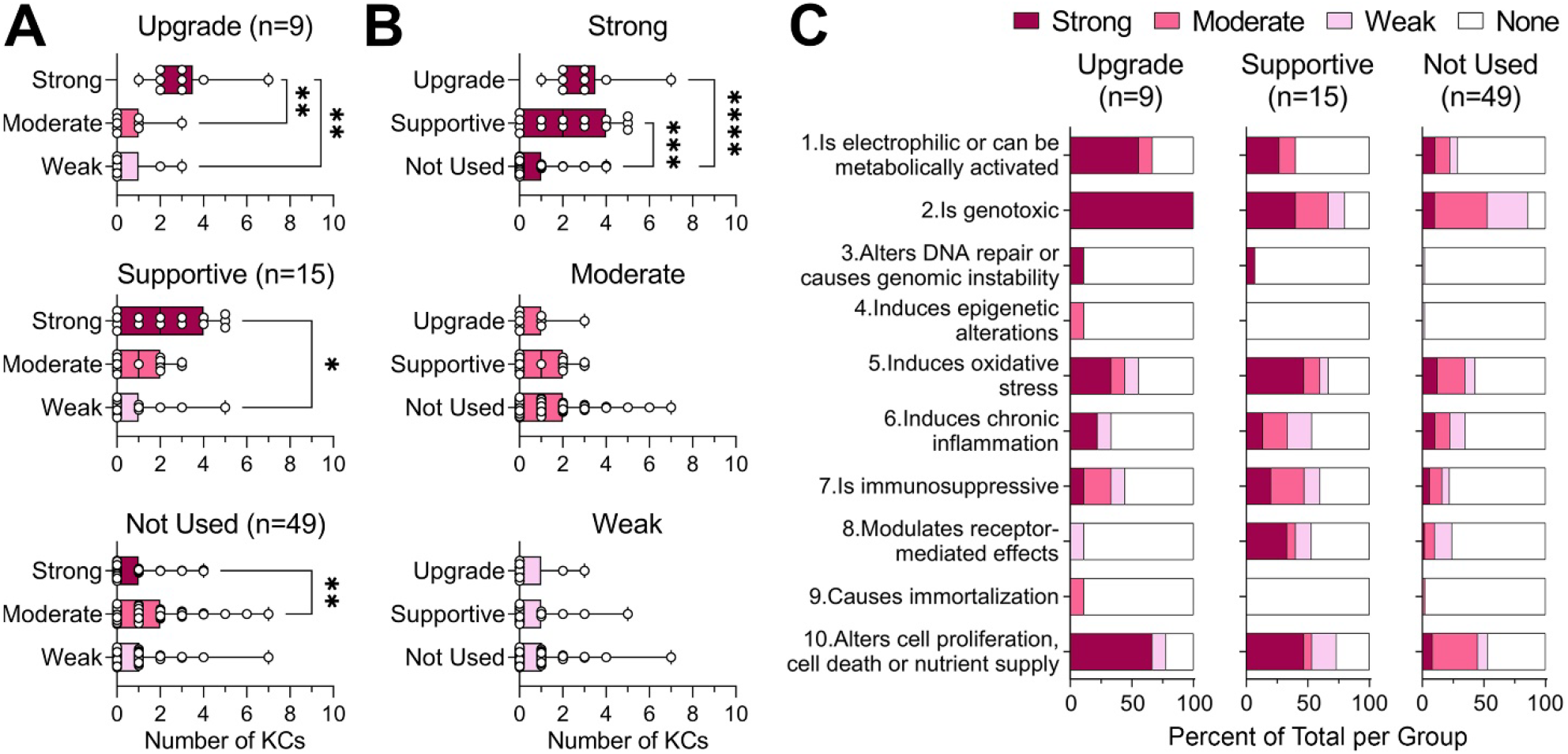
Mechanistic Evidence and strength of evidence for Key Characteristics used in IARC Monographs 112-130. (A) For each level of Mechanistic Evidence, number of Key Characteristics with each of the various levels is shown. (B) For each level of Key Characteristics, number of agents with each Mechanistic Evidence level shown. (C) Inverted stacked barplots displaying the strength of evidence for each Key Characteristic, with columns displaying the level of Mechanistic Evidence.

### Types of data used for Key Characteristics determinations by IARC Monographs Working Groups

Cancer hazard evaluations typically integrate evidence across several data types – evidence from exposed humans, human cell-based studies, studies in animals or animal-derived cells, as well as other model organisms (fish, invertebrates and procaryotes). In the past decade, data from many new approach methodologies, such as large-scale government toxicity screening efforts ToxCast/Tox21 (Dix et al. 2007; Richard et al. 2021; Thomas et al. 2019), have also been incorporated in a systematic manner into IARC Monographs process (Chiu et al. 2018). Figure 5 shows the types of evidence that was available for each Key Characteristic and whether studies were largely cohesive, equivocal, or showing lack of evidence for a particular key characteristic. The analysis by Krewski et al (2019a) concluded that for 86 known human carcinogens evaluated by IARC up to Monograph 106, evidence from exposed humans was abundant for four Key Characteristics – “1.Is electrophilic or can be metabolically activated,” “2.Is genotoxic,” “4.Induces epigenetic alterations,” and “6.Induces chronic inflammation.” However, agents that were evaluated in Monographs 112 through 130 had little data from exposed humans across all 10 Key Characteristics. Instead, information from the animal studies *in vivo* contributed the most to those cases where an affirmative link between an agent and a Key Characteristic was evident. The latter was especially evident for “10.Alters cell proliferation, cell death or nutrient supply” Key Characteristic. Conflicting results (labeled as equivocal in Figures 5 and 6) were most often present for “2.Is genotoxic” Key Characteristic. Interestingly, ToxCast data were most frequently used to refute a link between an agent and a Key Characteristic.

**Figure 5.**
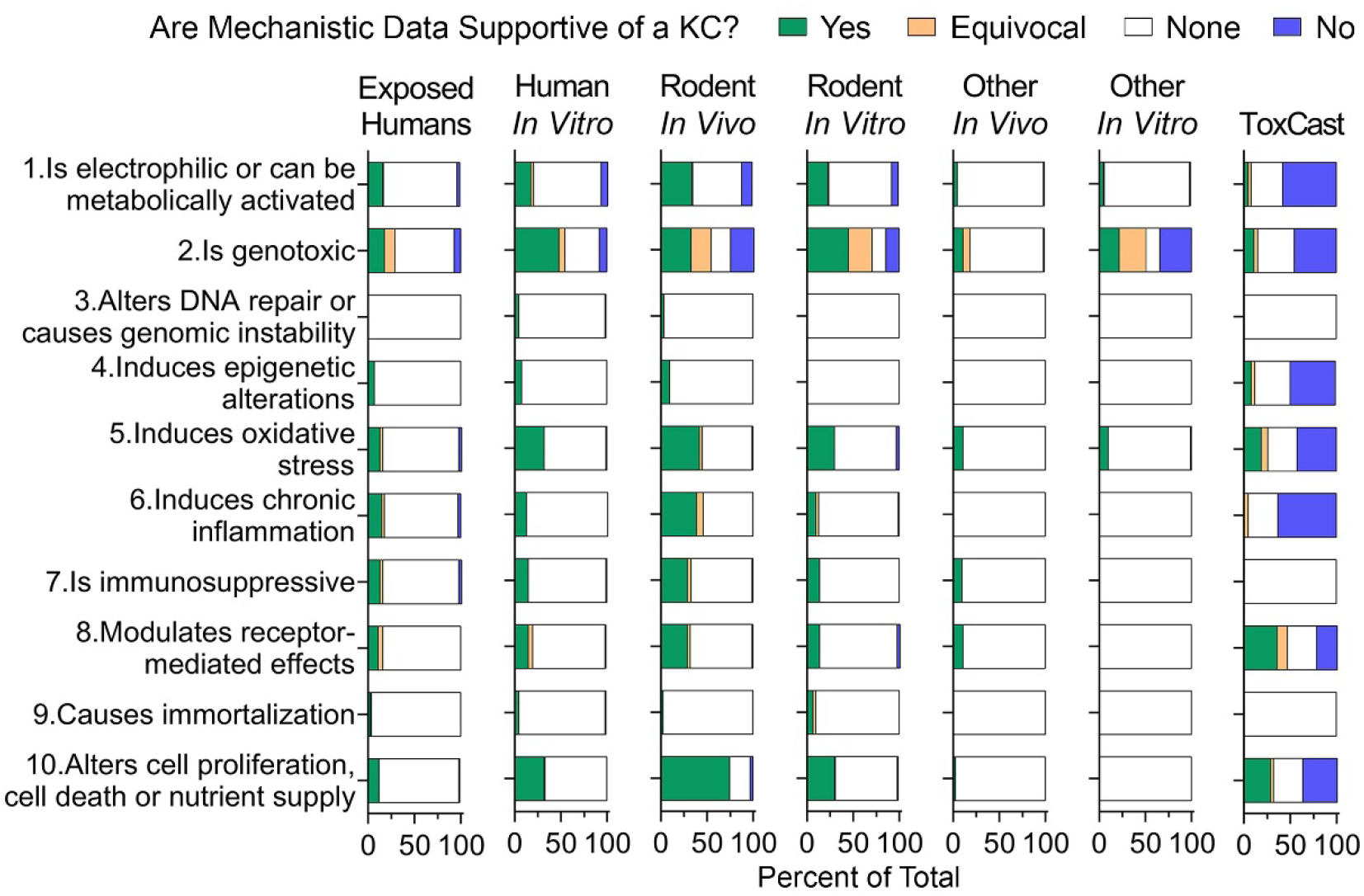
Inverted stacked barplots displaying the percentage of agents for each strength of Mechanistic Evidence level for each Key Characteristic and Model System.

**Figure 6.**
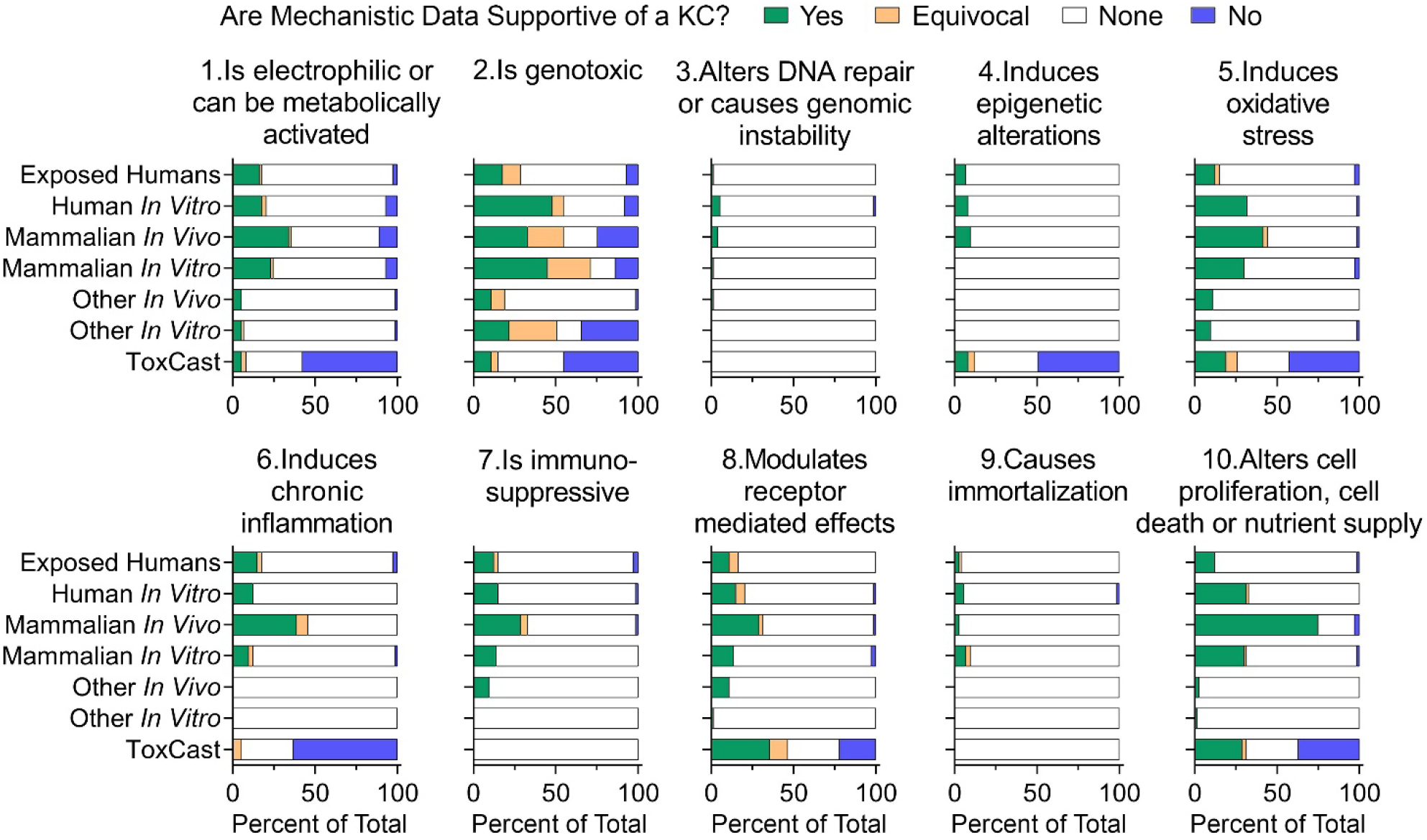
Inverted stacked barplots displaying the percentage of agents for each strength of Mechanistic Evidence level for each Model System and Key Characteristic.

When the same data as in Figure 5 are plotted for each Key Characteristic individually (Figure 6), additional patterns are more evident. While “2.Is genotoxic” Key Characteristic is most data-rich, it is noteworthy that it is data from human and animal *in vitro* models, followed by human and animal *in vivo* studies, not procaryotes, that has been most informative. Even though data from exposed humans is scarce overall, this data type contributed approximately evenly to 7 out of 10 Key Characteristics. Several Key Characteristics are very data-poor, indicative of either a lack of accessible assays to evaluate them either *in vivo* or *in vitro* (“3.Alters DNA repair or causes genomic instability” and “9.Causes immortalization”) or a mechanistic area that has only recently become accessible for study using sequencing techniques (“4.Induces epigenetic alteration”). Rapid expansion of evidence base for the latter Key Characteristic has been recently documented (Chappell et al. 2016; Goodman et al. 2022).

### Analysis of patterns in the use of Key Characteristics by IARC Monographs Working Groups

Because the analysis of mechanistic and other evidence by IARC Worksing Groups was done in each Monograph largely independently and more than 100 experts from over a dozen countries and many different areas of expertise participated in this process, it is interesting to determine whether any patterns emerge in co-occurrence of Key Characteristics, data types that may be most impactful, and the Key Characteristics that may have been most informative for the mechanistic upgrades. Because cancer is a disease with complex ethiology and multiple mechanisms operating simultaneously (Hanahan and Weinberg 2011; Pearce et al. 2015), patters would be expected. For this analysis, a statistical approach was utilized as detailed in Methods. In short, paiwise associations within and across data types and Key Characteristics and their application were examined as shown in Figure 1.

Because most agent-Key Characteristic pairs had no data for confident determination, the overall data matrix for this analysis is quite sparse. Two areas where mising information to preclude meaningful association analyses is least impactful were the “overall evaluation” determinations (strong, moderate, or weak) for each Key Characteristic [Figure 1, analysis type (I)], and cohesion and directionality determinations (yes, equivocal, or no) for ToxCast data [Figure 1, analysis type (II)]. Both datasets wer analysed using a categorical trend test with False Discovery Rate-based determination of significant associations. Figure 7A shows the results of this analysis for comparison (I), associations of the “overall strength” determination among Key Characteristics. Out of 45 possible pair-wise comparisons, only 3 were significant (*q*-value < 0.05). Key Characteristic “1.Is electrophilic or can be metabolically activated” was co-occurring with “2.Is genotoxic” and “10.Alters cell proliferation, cell death or nutrient supply.” In addition, Key Characteristic “2.Is genotoxic” was co-occurring with “5.Induces oxidative stress.” Figure 7B shows similar analysis for ToxCast data; out of possible 21 pair-wise comparisons (three Key Characteristics did not map to any ToxCast assays (Chiu et al. 2018)), 7 were significant. Interestingly, 4 out of these associations were driven mostly by the preponderance of the “no” determinations. The other three that were significantly concordant among themselves (“5.Induces oxidative stress,” “8.Modulates receptor-mediated effects,” and “10.Alters cell proliferation, cell death or nutrient supply”) had more balanced “yes” and “no” determinations that were concordant across agents (Supplemental Figure 1).

**Figure 7.**
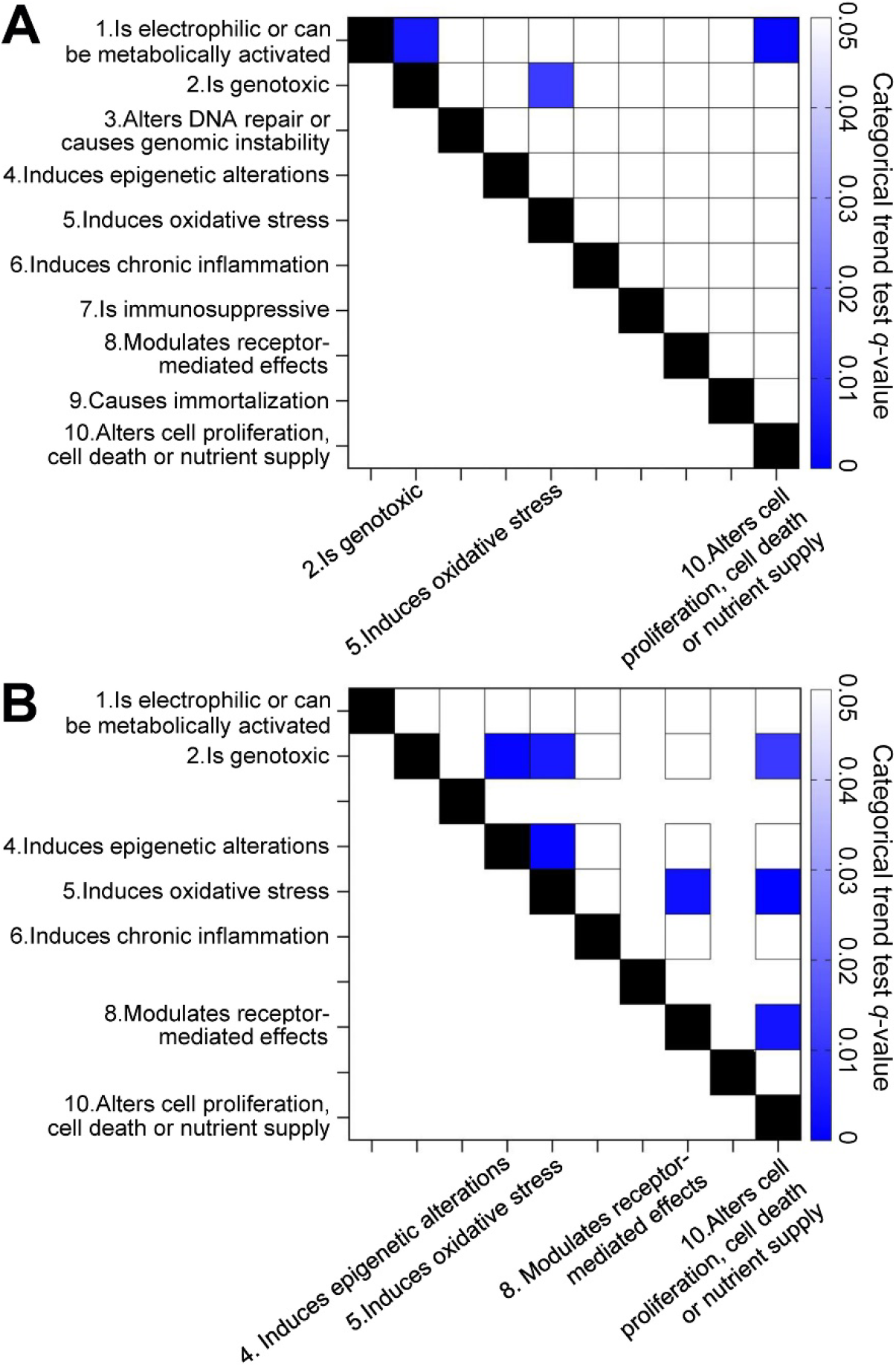
False discovery *q*-values for trend test pairwise comparisons of Key Characteristics. (A) Associations of the “overall strength” determination. (B) Associations of the ToxCast-mapped Key Characteristic data.

Concordance and directionality of the effect (yes, equivocal, or no) for each data type (“model system”) was used as a predictor of a dichotomized “overall strength” determination (strong vs moderate or weak, strong or moderate vs weak, strong vs weak, or moderate vs weak) for each Key Characteristic (Figure 1, analysis type (III)). Significance was determined using the Fisher’s exact test with Benjamini-Hochberg False Discovery Rate-based multiple testing correction. Remarkably, only one pair-wise association, between “2.Is genotoxic” Key Characteristic and Mammalian *in vitro* data type was significant when either strong or moderate determinations were compared against the weak ones (Figure 8A), or when strong and weak determinations were the only two options (Figure 8B).

**Figure 8.**
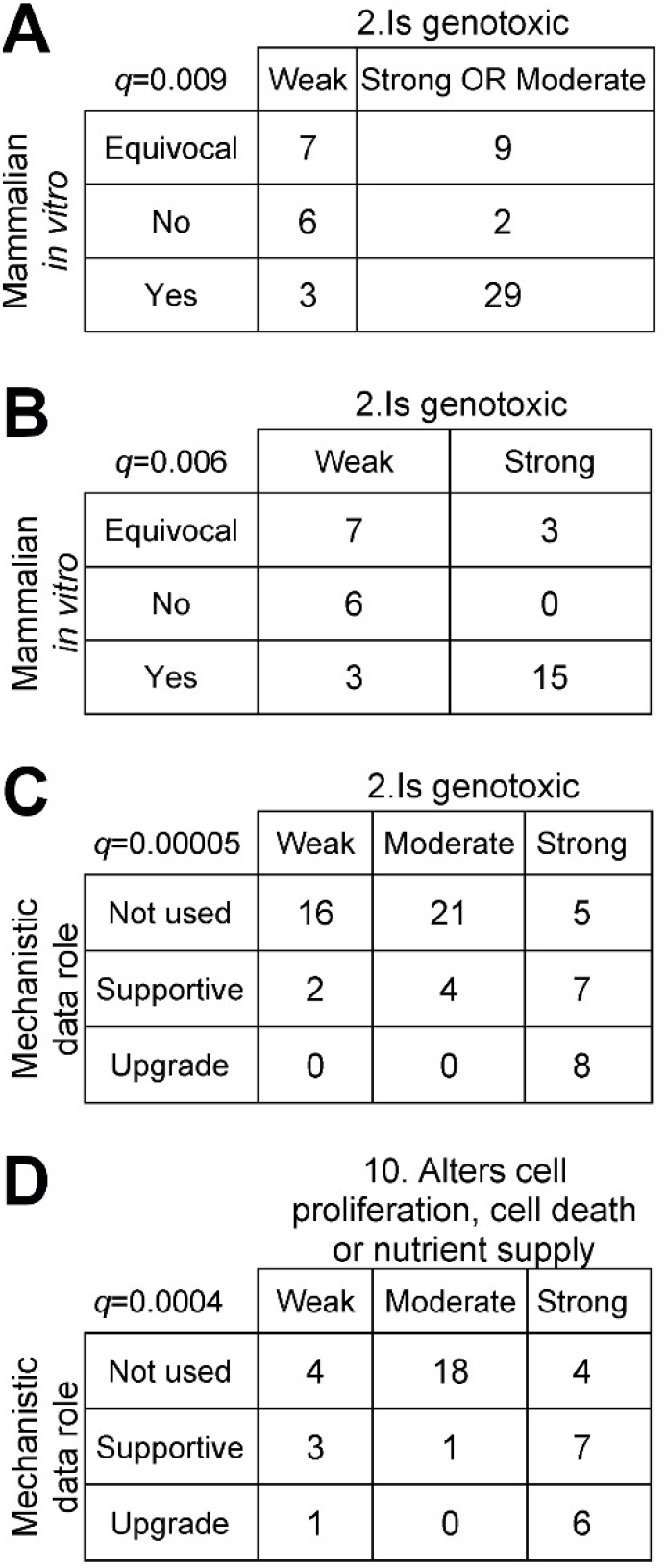
Comparison tables of strength of Mammalian *in vitro* data type shows significant associations (false discovery q<0.05) with Key Characteristic "2.is genotoxic" whether (A) the Key Characteristic value for Strong is combined with Moderate or (B) only Weak vs. Strong is used. (C) The Mechanistic data role shows a significant association with Key Characteristic "2.is genotoxic," as does (D) Key Characteristic "10.Alters cell proliferation, cell death, or nutrient supply."

Finally, the association of the “overall strength” (strong, moderate, or weak) of each Key Characteristic with the role that mechanistic data played in the overall cancer hazard classification (upgrade, supportive, or not used) was determined using the same type of analysis detailed above for Figures 8A-B (Figure 1, analysis type (IV)). The only significant results (Figures 8C and 8D) were found for the “2.Is genotoxic” and “10. Alters cell proliferation, cell death or nutrient supply” Key Characteristics. For Key Characteristic 2, weak determination of this Key Characteristic was most frequently co-occurring with mechanistic data not used for the final classification. Interestingly, while all instances of an upgrade that also had data for this Key Characteristic were when the evidence was deemed to be strong, comparable number of times a strong determination for this Key Characteristic occurred in cases when evidence was either supportive, or not used. The results were similar for Key Characteristic 10, and for both significant findings there was a clear trend that as the evidence for the Key Characteristic increased, the evidence for the Mechanistic data role also increased.

## Discussion

Virtually all regulatory decision frameworks for evaluating the potential of an agent to cause cancer in humans rely on observational epidemiological studies in humans, and/or experimental studies in rodents where tumor incidence is evaluated in all tissues and organs after long-term (typically 2 years or longer) exposures. Knowledge of the mechanisms of how the agent and disease (i.e., cancer) may be causally linked is always desirable, but in isolation (i.e., without associated epidemiological or animal studies) these data were not formally “actionable” for cancer hazard identification until recent amendments to the IARC Monographs Preamble (Samet et al. 2020). In spite of the major advances in formalizing the process of how mechanistic and other information is to be used by the regulators (Boobis et al. 2006; Meek et al. 2003; Sonich-Mullin et al. 2001), and its adoption into the official guidelines for carcinogen risk assessment (U.S. Environmental Protection Agency 2005), mechanistic data is almost always used only as supporting evidence. For example, (Thompson et al. 2011) noted that the mode of action analysis has historically been used to (i) judge the likelihood that adverse outcomes observed in animals will occur in humans, (ii) support the choice of low-dose extrapolation approaches, and/or (iii) identify data gaps that can be used to design additional studies to verify or refute the hypothesized key events. It is not uncommon, however, to find that the same overall “process” for cancer hazard identification and evidence database may result in opposing conclusions by the government agencies and others (Cohen et al. 2004; Felter et al. 2018; Klaunig et al. 2003; Portier et al. 2016).

Indeed, molecular, cellular, organ- or species-specific mechanistic events that may be involved in carcinogenesis are complex, and their interpretation requires subject matter expertise. Concerns about possible implicit bias of the experts who are tasked with evaluation of the mode of action data in course of cancer hazard identification are likely to be no different from concerns about bias in decisions by other health professionals (FitzGerald and Hurst 2017). In addition, the practice of systematic literature searching to identify relevant mechanistic evidence to facilitate weight of evidence analyses (Weed 2005) has not been commonplace until about a decade ago, generally because of the large number and variety of studies. After the evidence has been identified, the challenge of “weighting” diverse and voluminous data is amplified if the assessments are performed by *ad hoc* committees of experts, such as IARC Monographs Working Group members. Indeed, a systematic re-evaluation of all Group 1 human carcinogens by IARC identified several challenges that needed to be addressed to increase consistency, minimize implicit bias, and ensure comprehensive nature of evaluations (Birkett et al. 2019; Smith et al. 2016). These challenges included the recognition of both diversity and complexity of the mechanisms that may be pertinent to each agent, rapid evolution of the new methods (e.g., - omics) and models (e.g., transgenic animals and complex cell-based constructs) to study the mechanisms, and the need to develop systematic search methods for complex mechanistic events that may lead to cancer.

The IARC’s approach to address these challenges is known as the Key Characteristics of Carcinogens (Smith et al. 2016); it was applied both retroactively to all Group 1 agents (Krewski et al. 2019a), as well as used in all IARC Monographs since 2015 (Guyton et al. 2015; Guyton et al. 2018a). Previous publications concluded that application of the Key Characteristics approach in cancer hazard identification, combined with transparent documentation of both evidence and decisions (Shapiro et al. 2018), is very useful for organizing and evaluating the mechanistic data because it enables a reproducible systematic review of the mechanistic evidence and avoids the need to identify *a priori* specific pathways and hypotheses. It was also noted that because the Key Characteristics have been derived by a diverse group of experts in toxicology, epidemiology, pathology, and risk assessment (Al-Zoughool et al. 2019; Birkett et al. 2019; Smith et al. 2016), they are rooted in comprehensive empirical evidence of the mechanisms associated with diverse (physical, chemical and biological) known carcinogens and thus provide an agnostic and unbiased pathway for evaluation of other agents. Not only has this approach been used by IARC Monographs to evaluate more than 73 agents (through Monograph 130), it also has been used in cancer hazard identification for several dozen agents by US EPA Integrated Risk Information System (IRIS) and the National Toxicology Program’s Report on Carcinogens (RoC) (Supplemental Table 5). In addition to the IARC Preamble (IARC Monographs Programme 2019), there are detailed instructions and search terms that explain how the Key Characteristics approach is applied by both IRIS and RoC programs (NTP 2022; US EPA 2022).

Our analysis of the use of Key Characteristics and their application in the cancer hazard conclusions by 19 IARC Monograph Working Groups spans information from a 7-year period (2015-2022) and is informative in several ways. First, we found that for most Key Characteristics and agents there was little data (∼66% of the instances). Even when the data was available, the Working Group members deemed any Key Characteristic as “strong” for only ∼12.5% of the instances; there were on average 1.3 “strong” Key Characteristics per agent. As found previously for 86 Group 1 agents (Krewski et al. 2019a), or 34 agents classified into Groups 1, 2A or 2B (Guyton et al. 2018a), there is a wide range in the number of strong, moderate, or weak Key Characteristic determinations per agent, irrespective to which cancer hazard group they were ultimately classified. Still, there is a significant trend in the number of strong Key Characteristics among groups – Group 1 agents on average have the most, even though the ranges are overlapping. Collectively, these observations show that IARC Monographs Working Group members are very cautious in weighing the strength of mechanistic evidence for each agent and each Key Characteristic.

Second, while it was not surprising that Key Characteristics with the most determinations available correspond to well-known mechanisms of carcinogenesis (i.e., genotoxicity, cell death/ proliferation, oxidative stress, metabolic activation, immune-mediated events, chronic inflammation, and receptor-mediated events), it was noteworthy that some well-established cancer mechanisms had very little data. For example, it is well established that induction of DNA repair (Powell et al. 2005) and associated genome instability (Hanahan and Weinberg 2011) are enabling carcinogenic transformation of cells; however, few studies evaluate these endpoints in the context of mechanistic study of environmental agents. Similarly, epigenetic alterations are well-recognized as chemical carcinogen-associated mechanisms (Herceg et al. 2013; Miller 1970), yet the molecular tools to study these have only become widely available in the last 10-15 years which led to a rapid increase in studies of epigenetic effects of chemicals (Chappell et al. 2016; Goodman et al. 2022). Finally, replicative immortality is one of the originally identified hallmarks of cancer cells (Hanahan and Weinberg 2011), yet it is a process that is difficult to evaluate in traditional animal or cell-based studies of chemical exposures.

Third, our analysis showed that not only are there differences in the amount and type of evidence for different Key Characteristics, but also that mechanistic data from *in vivo* studies in animals was the greatest contributor to the associations across all Key Characteristics. Studies in human and rodent cells *in vitro* were also informative and more abundant than mechanistic evidence from “exposed humans.” This finding is instructive in the context of the increasing shift to *in vitro* methods in drug and chemical safety evaluations (Anklam et al. 2022; Editorial 2013). Notably, *in vitro* data from ToxCast assays, a most comprehensive source of non-animal evidence because thousands of chemicals were tested in hundreds of assays and the data are publicly available (Williams et al. 2017), contributed little as supportive evidence for most agents and seven Key Characteristics that could be mapped to ToxCast (Chiu et al. 2018). Instead, ToxCast data were most informative in conclusions that a Key Characteristic was not associated with an agent. This observation is important because it provides confidence in the comprehensive nature of the overall evidence base and that few “blind spots” may exist in terms of potential mechanisms of carcinogenesis for the agent under consideration. Still, the observation that members of the IARC Monographs Working Groups were more acceptable of the mechanistic data from whole animal experiments, rather than cell-based studies, shows the challenging path forward for decision-making on the agents that lack human or animal data.

Fourth, concerns have been voiced that decisions about cancer hazard could be made on certain Key Characteristics in isolation, for example “5.Induces oxidative stress” (Bus 2017). While this is a potentially valid concern, also recognized in the Preamble (IARC Monographs Programme 2019), there was no instance when such isolated determination was made. Both retrospective analysis of Group 1 agents (Krewski et al. 2019a) and the analysis of the application of the Key Characteristics since 2015 [(Guyton et al. 2018a) and this manuscript] have shown that multiple Key Characteristics are typically deemed as strong among the agents classified into Group 1. In fact, we found that in the instances where a mechanistic “upgrade” was considered, it was “2.Is genotoxic” and “10.Alters cell proliferation, cell death or nutrient supply” that showed a significant association with a Working Group’s decision – either to upgrade, to use mechanistic data as supportive evidence, or not use these data at all. If anything, it was weak or moderate evidence in these two Key Characteristics that determined the decisions to not use the mechanistic data in the final decision, and not that strong evidence tipped the scales. Similarly, these two Key Characteristics were strongly associated with each other across all evaluations – both significantly co-occurred with the “1.Is electrophilic or can be metabolically activated” Key Characteristic which is a known precursor event and a mechanism that is difficult to study in cell-based models (Ooka et al. 2020).

Finally, it is noteworthy that we found few significant co-occurrence associations among Key Characteristics and between Key Characteristics and ultimate decisions on what role mechanistic evidence had in the final cancer hazard classification. The finding may inform potential future attempts to substitute expert judgement with artificial intelligence approaches. The paucity of discernable patterns across a large database of 73 agents is instructive insofar as the expert judgement and group discussions may be difficult to replace with computational algorithms. While efforts to automate the systematic literature review process, including search, screening, and data extraction have enjoyed several advances and software tools are available and used (Marshall and Wallace 2019; van Dinter et al. 2021), the process of weighing and synthesizing evidence is still the domain of expert judgement and subject to external peer review (National Toxicology Program 2019; US EPA 2022). Indeed, interpretation of the data for each individual Key Characteristic, and integrating evidence across all 10, are processes that involve intense discussions and debate among experts consider the mechanistic evidence and how it may support or refute other evidence streams (i.e., human and animal evidence). The Key Characteristics are not used by IARC Working Groups to predict cancer (Becker et al. 2017) or to determine carcinogenic potency (Goodman et al. 2018; Guyton et al. 2018b).

Overall, this study aimed to demonstrate how the Key Characteristics approach has been implemented by IARC Monographs Working Groups since 2015 and to address some of the concerns with the approach (Becker et al. 2017; Bus 2017; Goodman and Lynch 2017; Goodman et al. 2018; Meek and Wikoff 2023). In the past decade, the Key Characteristics have been used in cancer hazard evaluations of over 100 agents by IARC, US EPA and NTP RoC programs, thus this type of a mechanistic construct has enjoyed wide, but not unanimous adoption by the decision-makers. Additional Key Characteristics for chemical-induced repeat dose toxicity have been developed for other organs (Arzuaga et al. 2019; Germolec et al. 2022; Jennings et al. 2023; La Merrill et al. 2020; Lind et al. 2021; Rusyn et al. 2021); the overall approach is gaining recognition and acceptance both in the US and Europe. Through the analysis of the patterns and associations in the application of the Key Characteristics, the field of regulatory science can learn from the contexts of how and when the mechanistic data can be integrated into decisions.

Among the key findings is the observation that there the gaps in available data for Key Characteristics for many agents. This situation is even more dire for thousands of agents that have had no formal evaluations of the cancer hazard potential. For positive association decisions, the Working Group experts relied heavily on the mechanistic *in vivo* animal data that are available for several Key Characteristics, *in vitro* data from animal and human models was also used. To exclude the involvement of the Key Characteristics, the ToxCast data were most informative. In fact, available human *in vitro* data were much more likely to be interpreted as supportive of a Key Characteristic than the ToxCast data. Although further investigation is warranted, this finding may be suggestive of greater sensitivity of newer human *in vitro* model systems to the underlying mechanisms of carcinogenesis. As human *in vitro* new approach methodologies continue to develop, the potential for high-throughput organotypic assays might be viewed in this light. Overall, the increased availability of systemized data streams, such as human *in vitro* data, would provide the basis for confident conclusions about both positive and negative associations and constitute relative contributions of various sources of weight in expert judgments.

## Acknowledgements

This work was supported, in part, by grants from the National Institute of Environmental Health Sciences (P42 ES027704) and the U.S. Environmental Protection Agency (RD84045001). The views expressed in this manuscript do not reflect those of the funding agencies. The use of specific commercial products in this work does not constitute endorsement by the funding agencies.

