## Supplemental Figure 1 for "Ten Years of Using Key Characteristics of Human Carcinogens to Organize and Evaluate Mechanistic Evidence in IARC Monographs on the Identification of Carcinogenic Hazards to Humans: Patterns and Associations"

**Supplemental Figure 1.** Associations of ToxCast data across different Key Characteristics. (A) Associations that were driven largely by the negative ToxCast data. (B) Associations that were driven largely by both negative and positive ToxCast data. The association in the ToxCast data determinations for each pairwise KC combination [e.g., KC1 🡪 KC2] were examined from a total of 45 possible comparisons. The analysis was conducted using Fisher’s exact test with False Discovery Rate (FDR) – based testing to determine significant (*q* < 0.05) associations.

**(A)**

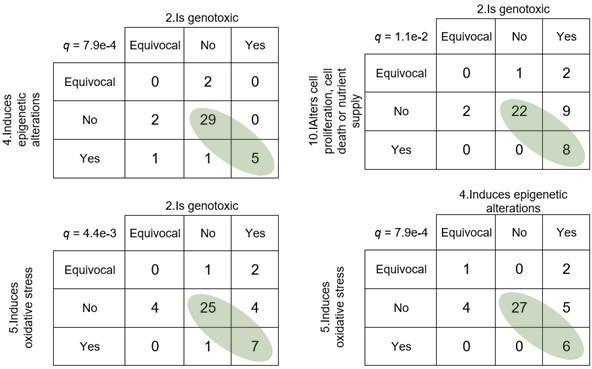

**(B)**

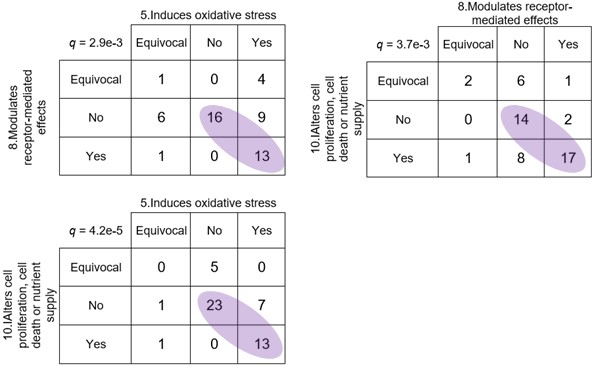
