## Supplemental Table 1 for "Ten Years of Using Key Characteristics of Human Carcinogens to Organize and Evaluate Mechanistic Evidence in IARC Monographs on the Identification of Carcinogenic Hazards to Humans: Patterns and Associations"

**Supplemental Table 1.** IARC Monographs on the Identification of Carcinogenic Hazards to Humans publications included in this study.

| **Substance/Dietary factor/Occupation** | **Volume [Year]** | **URL for the Monographs** |
| --- | --- | --- |
| - IARC Monographs Preamble – Preamble to the IARC Monographs (amended January 2019) | 2019 | <https://monographs.iarc.who.int/iarc-monographs-preamble-preamble-to-the-iarc-monographs/> |
| - Malathion - Parathion - Diazinon - Glyphosate - Tetrachlorvinphos | 112  [2015] | <https://publications.iarc.fr/549> |
| - DDT - Lindane - 2,4-D | 113  [2015] | <https://publications.iarc.fr/550> |
| - Red and processed meat | 114  [2015] | <https://publications.iarc.fr/564> |
| - 1-Bromopropane - 2-Metcaptobenzothiazole - 3-chloro-2-methylpropene - N,N-Dimethylformamide - N,N-Dimethyl-p-toluidine - Hydrazine - Tetrabromobisphenol A | 115  [2016] | <https://publications.iarc.fr/563> |
| - Drinking Coffee - Drinking mate and very hot beverages | 116  [2016] | <https://publications.iarc.fr/566> |
| - Pentachlorophenol - 2,4,6-Trichlorophenol - 3,3ʹ,4,4ʹ-Tetrachloroazobenzene - Aldrin - Dieldrin | 117  [2016] | <https://publications.iarc.fr/574> |
| - Welding - Molybdenum trioxide - Indium Tin Oxide | 118  [2017] | <https://publications.iarc.fr/569> |
| - 1-Tert-Butoxypropan-2-ol - b-Myrcene - Furfuryl alcohol - Melamine - Pyridine - Tetrahydrofuran - Vinylidene chloride | 119  [2017] | <https://publications.iarc.fr/575> |
| - Benzene | 120  [2017] | <https://publications.iarc.fr/576> |
| - Styrene - Styrene-7,8-oxide - Quinoline | 121  [2018] | <https://publications.iarc.fr/582> |
| - Isobutyl Nitrate - b-picoline - Methyl acrylate - Ethyl acrylate - 2-ethylhexyl acrylate - Trimethylolpropane triacrylate | 122  [2018] | <https://publications.iarc.fr/583> |
| - 2-Chloronitrobenzene - 4- Chloronitrobenzene - 1,4-Dichloro-2-nitobenzene - 2,4- Dichloro-1-nitobenzene - 2-Amino-4-chlorophenol - Ortho-phenylenediamine & ortho-phenylenediamine dihydrochloride - Para-nitroanisole - N,N-Dimethylacetamide | 123  [2018] | <https://publications.iarc.fr/584> |
| - Night shift work | 124  [2019] | <https://publications.iarc.fr/593> |
| - Allyl chloride - 1-Bromo-3-chloropropane - 1-Butyl glycidyl ether - 4-Chlorobenzotrifluoride - Glycidyl methacrylate | 125  [2019] | <https://publications.iarc.fr/596> |
| - Opium consumption | 126  [2020] | <https://publications.iarc.fr/600> |
| - Ortho-anisidine & ortho-anisidine hydrochloride - Ortho-nitroanisole - Aniline and aniline hydrochloride - Cupferron | 127  [2020] | <https://publications.iarc.fr/599> |
| - Acrolein - Crotonaldehyde - Arecoline | 128  [2020] | <https://publications.iarc.fr/602> |
| - Gentian violet - Malachite green - CI direct blue 218 | 129  [2021] | <https://publications.iarc.fr/603> |
| - 1,1,1-Trichloroethane - 1,2-Diphenylhydrazine - Diphenylamine - N-Methylolacrylamide - Isophorone | 130  [2021] | <https://publications.iarc.fr/611> |
