## Supplemental Table 5 for "Ten Years of Using Key Characteristics of Human Carcinogens to Organize and Evaluate Mechanistic Evidence in IARC Monographs on the Identification of Carcinogenic Hazards to Humans: Patterns and Associations"

**Supplemental Table 5.** Cancer hazard identification assessments by US EPA Integrated Risk Information System (IRIS) and the National Toxicology Program’s Report on Carcinogens (RoC) that used Key Characteristics approach to evaluate mechanistic evidence.

5A. US EPA evaluations.

| **Substance** | **Year** | **URL for the Assessment Scoping Documents** |
| --- | --- | --- |
| ORD Staff Handbook for Developing IRIS Assessments | 2022 | [https://ordspub.epa.gov/ords/eims/eimscomm.getfile?p_download_id=545991](https://urldefense.com/v3/__https:/ordspub.epa.gov/ords/eims/eimscomm.getfile?p_download_id=545991__;!!KwNVnqRv!B5UVRY17dwe9DFZIZpk5svZ6XeXnhjwEsDJoJOmsNPm2nqeVJtMffGgPUmMnWhlZVToDYQbFwLxAVeQqHc7C$) |
| - PFBA (375-22-4) - PFHxA (307-24-4) - PFHxS (355-46-4) - PFNA (375-95-1) - PFDA (335-76-2) | 2021 | <https://ordspub.epa.gov/ords/eims/eimscomm.getfile?p_download_id=542033> |
| - Cobalt and Cobalt Compounds | 2022 | <https://ordspub.epa.gov/ords/eims/eimscomm.getfile?p_download_id=545669> |
| - Hexavalent Chromium [Cr(VI)] | 2022 | <https://ordspub.epa.gov/ords/eims/eimscomm.getfile?p_download_id=545542> |
| - Naphthalene | 2023 | <https://ordspub.epa.gov/ords/eims/eimscomm.getfile?p_download_id=546419> |
| - Ethylbenzene | 2023 | [https://ordspub.epa.gov/ords/eims/eimscomm.getfile?p_download_id=546289](https://urldefense.com/v3/__https:/ordspub.epa.gov/ords/eims/eimscomm.getfile?p_download_id=546289__;!!KwNVnqRv!B5UVRY17dwe9DFZIZpk5svZ6XeXnhjwEsDJoJOmsNPm2nqeVJtMffGgPUmMnWhlZVToDYQbFwLxAVYyToXa2$) |

5B. NTP RoC evaluations.

| **Substance** | **Year** | **URL for the Monographs or Ongoing Assessments** |
| --- | --- | --- |
| Standard Search Strings for Literature Database Searches: Appendix to the Handbook for Preparing Report on Carcinogens Monographs | 2022 | <https://ntp.niehs.nih.gov/ntp/roc/handbook/rochandbookappendix_508.pdf> |
| - Antimony Trioxide | 2018 | <https://ntp.niehs.nih.gov/ntp/roc/monographs/antimony_final20181019_508.pdf> |
| - Bromochloroacetic acid - Bromodichloroacetic acid - Chlorodibromoacetic acid - Dibromoacetic acid - Dichloroacetic acid - Tribromoacetic acid | 2018 | <https://ntp.niehs.nih.gov/ntp/roc/monographs/haafinal_508.pdf> |
| - Halogenated Flame Retardants | Ongoing | <https://ntp.niehs.nih.gov/whatwestudy/assessments/cancer/ongoing/hfr> |
| - Selected Nitropolycyclic Aromatic Hydrocarbon Compounds | Ongoing | <https://ntp.niehs.nih.gov/whatwestudy/assessments/cancer/ongoing/npah> |
| - Wood Smoke | Ongoing | <https://ntp.niehs.nih.gov/whatwestudy/assessments/cancer/ongoing/woodsmoke> |
| - Para-Chlorotrifluorotoluene | Ongoing | <https://ntp.niehs.nih.gov/whatwestudy/assessments/cancer/ongoing/pctft> |
| - Polycyclic Aromatic Hydrocarbons (PAHs) | Ongoing | <https://ntp.niehs.nih.gov/whatwestudy/assessments/cancer/ongoing/pahs> |
